# Selective sweeps influence diversity over large regions of the mouse genome

**DOI:** 10.1101/2021.06.10.447924

**Authors:** Tom R. Booker, Benjamin C. Jackson, Rory J. Craig, Brian Charlesworth, Peter D. Keightley

## Abstract

To what extent do substitutions in protein-coding versus gene-regulatory regions contribute to fitness change over time? Answering this question requires estimates of the extent of selection acting on beneficial mutations in the two classes of sites. New mutations that have advantageous or deleterious fitness effects can induce selective sweeps and background selection, respectively, causing variation in the level of neutral genetic diversity along the genome. In this study, we analyse the profiles of genetic variability around protein-coding and regulatory elements in the genomes of wild mice to estimate the parameters of positive selection. We find patterns of diversity consistent with the effects of selection at linked sites, which are similar across mouse taxa, despite differences in effective population size and demographic history. By fitting a model that combines the effects of selective sweeps and background selection, we estimate the strength of positive selection and the frequency of advantageous mutations. We find that strong positive selection is required to explain variation in genetic diversity across the murid genome. In particular, we estimate that beneficial mutations in protein-coding regions have stronger effects on fitness than do mutations in gene-regulatory regions, but that mutations in gene-regulatory regions are more common. Overall though, our parameter estimates suggest that the cumulative fitness changes brought about by beneficial mutations in protein-coding may be greater than those in gene-regulatory elements.

## Introductions

Understanding the relative contributions of protein-coding and gene regulatory variation to adaptation is a long-standing goal of evolutionary biology. Molecular changes in protein-coding and gene regulatory regions contribute to evolution, but in classic essays King and Wilson (1975) and Carroll (2005) argued that changes in gene expression may dominate adaptive evolution. King and Wilson (1975) reasoned that since nucleotide identity between human and chimpanzee proteins is around 99%, there are too few protein sequence difference between the species, implying that changes in gene regulation are probably required to explain the many phenotypic differences between the species. Carroll’s (2005) argument highlighted the idea that molecular changes in the gene regulatory apparatus may have smaller pleiotropic effects than those in protein-coding regions, so that changes in gene expression may dominate adaptive evolution. However, Hoekstra and Coyne (2007) attempted to refute these arguments, maintaining that there is insufficient evidence to decide whether adaptation is primarily driven by changes in protein sequences or gene regulatory elements. For example, a 1% difference in protein sequence between humans and chimpanzees could still result in a very large number of phenotypic differences. However, the contribution of individual variants to additive genetic variance for a trait is expected to be proportional to the square of their phenotypic effect sizes, assuming semi-dominance (Fisher 1918; Falconer and Mackay 1996). Without an understanding of the frequencies of new mutations, their effect sizes, and the strength of selection acting on them, the question of the contribution of molecular evolution in different genomic elements to adaptation will remain intractable.

Information on the strength of selection acting on beneficial mutations and the rates at which they occur can be obtained by analysing patterns of neutral genetic diversity. Because genetically linked sites do not evolve independently, natural selection acting at a given site may leave signatures at linked sites that are informative about the strength and mode of selection. The effects of selection at linked sites on neutral genetic diversity depend on the frequency and strength of selected mutations and the rate of recombination (Charlesworth 2012; Hermisson and Pennings 2017; Stephan 2019). Several modes of selection at linked sites have been identified. Of specific relevance to this study are background selection (BGS), caused by the removal of deleterious mutations from a population, and selective sweeps, caused by the spread of advantageous variants. The classic footprint of a selective sweep is a trough in nucleotide diversity at neutral sites surrounding an adaptive substitution. The reduction in nucleotide diversity caused by a sweep is proportional to the ratio of the strength of selection acting on the causal mutation to the local recombination rate (Barton 2000). Using such information, Wiehe and Stephan (1993) developed a model of recurrent selective sweeps and used it to estimate the frequency and strength of advantageous mutations in *Drosophila melanogaster*. They fitted their model of sweeps to the relationship between recombination rate and nucleotide diversity for a number of loci sampled across the *D. melanogaster* genome. At the time of their analysis, the theory of BGS was in its infancy, and models combining the effects of BGS and sweeps had not been developed. However, the effects of BGS are expected to be ubiquitous across the genome (McVicker et al. 2009; Comeron, 2014; Elyashiv et al. 2016; Pouyet et al. 2018), and conceptually similar studies to Wiehe and Stephan (1993) have shown that controlling for BGS is important when parametrizing sweep models (Kim and Stephan 2000; Comeron 2014; Elyashiv et al. 2016; Campos et al. 2017).

In *Drosophila*, there are reductions in average diversity around recent nonsynonymous substitutions, which are greater than those observed around synonymous substitutions (Sattath et al. 2011; Elyashiv et al. 2016). To investigate the causes of this difference, Elyashiv et al. (2016) fitted a model of sweeps and BGS to genome-wide variation in genetic diversity in *D. melanogaster* and found that a combination of BGS and selective sweeps provided a close fit to the observed data. From the fit of their model to empirical data, Elyashiv et al. (2016) inferred a distribution of fitness effects for advantageous mutations that included a class of very strongly selected mutations and a more mildly beneficial class. In both mice and humans, however, there is very little difference between the profiles of diversity around recent nonsynonymous and synonymous substitutions (Hernandez et al. 2011; Halligan et al. 2013). In these species, dips in average nucleotide diversity have been observed in genomic regions surrounding whole functional elements, such as protein-coding exons or conserved non-coding elements, which may reflect the cumulative effects of recurrent selective sweeps and BGS (Hernandez et al. 2011; Halligan et al. 2013; Booker and Keightley 2018)

Natural populations of mice in the genus *Mus* are excellent material for the study of adaptive evolution in different regions of the mammalian genome. Their populations are very large compared to other mammals (Leffler et al. 2012), so there is likely to be more power for population genetic analyses to differentiate between the evolutionary processes that affect genetic variability. Previous studies in mice have established that both protein-coding genes and regions putatively involved in gene regulation have an excess of sequence differences from sister taxa compared to that expected under a model of purifying selection, suggesting widespread adaptive molecular evolution (Halligan et al. 2010, 2013). Halligan et al. (2013) analysed a sample of *Mus musculus castaneus* individuals and estimated that there have been around 1.3 million and 0.38 million positively selected regulatory and nonsynonymous changes, respectively, over the period since this subspecies began to diverge from rats. At face value, this finding suggests that changes in gene regulation may dominate adaptive evolution in mice. However, Halligan et al. (2013) also showed that there are much larger reductions in neutral diversity surrounding protein-coding exons than around gene regulatory elements, and that BGS could not fully explain these observations (Halligan et al. 2013; Booker and Keightley 2018). Halligan et al. (2013) concluded that this difference in neutral diversity may reflect differences in the strength of positive selection acting on the different classes of sites.

Building on Halligan et al. (2013), we have sought to tease apart the contributions of BGS and sweeps to the patterns of nucleotide diversity observed in the Eastern house mouse *M. m. castaneus* (Booker and Keightley 2018). We inferred the distribution of fitness effects (DFE) for deleterious and advantageous mutations occurring in protein-coding genes and gene regulatory elements, by analysing the frequency distribution of derived allele frequencies (the unfolded site frequency spectrum, uSFS). Based on analysis of the uSFS, we found that a model of positive selection was insufficient to explain the troughs in nucleotide diversity around protein-coding exons or conserved non-coding elements (CNEs). However, we found that infrequent, strongly beneficial mutations, which have negligible effect on the uSFS, potentially could do so (Booker and Keightley 2018). This is because infrequent, strongly advantageous mutations may substantially influence diversity at linked sites, while making very little contribution to the uSFS. We concluded that the parameters of positive selection are very difficult to accurately estimate from the uSFS alone (Booker 2020). To understand the relative strengths of selection acting on protein-coding versus gene regulatory regions, the analysis of a model of selective sweeps fitted to patterns of neutral genetic variability may be more powerful.

In this study, we examine the reductions in nucleotide diversity surrounding protein-coding exons and conserved non-coding elements in wild mice, and attempt to tease apart the modes of selection operating on the two different elements. We fitted a model of selective sweeps to the patterns observed in *M. m. castaneus*, while correcting for the confounding effects of BGS. Our analysis provides evidence that the strength of selection acting on beneficial mutations in protein-coding exons is far greater than that acting on conserved non-coding elements. Using a simple model of the fitness change brought about by positive selection, we find that selection on protein-coding regions may contribute more to fitness change, despite positive selection occurring more frequently in regulatory regions of the genome. We then compared patterns of putatively neutral diversity among the principal subspecies of *Mus musculus* and their sister species *Mus spretus*. We find that the profiles of nucleotide diversity and the inferred distributions of fitness effects among each group are similar, suggesting that the contributions of positive selection to protein-coding and regulatory change are similar in the different mouse taxa. Note that our goal in this study is not to identify the individual loci that selection has recently acted on; for a recent study identifying the targets of recent selection in wild mice see (Lawal et al. 2021).

## Results and Discussion

### Profiles of genetic diversity around protein-coding exons and conserved non-coding elements in multiple mouse lineages

If different mouse lineages are subject to similar selection pressures, we might expect that they exhibit similar patterns of diversity across their genomes. We thus compared patterns of genetic diversity in populations of the house mouse *Mus musculus* and the sister species *Mus spretus*. We analysed data previously reported by Halligan et al. (2013) and Harr et al. (2016) for the two mouse species, *M. musculus* and *M. spretus*. For *M. musculus*, we analysed samples from the three sub-species, *M. m. castaneus, M. m. domesticus* and *M. m. musculus*. The *M. m. castaneus* individuals (*n* = 10) were from Himachal Pradesh, India. For *M. m. domesticus*, populations were sampled in France (*n* = 8), Germany (*n* = 8) and Iran (*n* = 8). In the case of *M. m. musculus*,populations were sampled in Afghanistan (*n* = 6), the Czech Republic (*n* = 8) and Kazakhstan (*n* = 8). The *M. spretus* individuals were sampled in Spain (*n* = 8). We refer to the different sub-populations of *M. m. domesticus* and *M. m. musculus* by the countries where the individuals were sampled.

We identified conserved non-coding elements (CNEs) in murid rodents using a 40-way alignment of placental mammals by means of the *phastCons* approach (Siepel et al. 2005). Following Williamson et al. (2014), the genomes of *M. musculus* and other rodents were masked in the alignment to limit ascertainment bias affecting elements that have recently diverged in the rodent lineage. CNEs identified using *phastCons* overlap with features such as promoters and enhancers (Lindblad-Toh et al. 2011), and thus are likely to have roles in the regulation of gene expression.

For each of the mouse taxa, we examined putatively neutral nucleotide diversity (*π*) in genomic regions surrounding protein-coding exons and CNEs using the methods described by Halligan et al. (2013). Briefly, polymorphism data were extracted in genomic windows surrounding protein-coding exons and CNEs. We masked any putatively functional sites from analysis windows; these included the exons (including UTRs) of genes annotated in the *M. musculus* genome by ENSEMBL in release 93 (Howe et al. 2021) and CNEs. For each analysis window, we calculated the genetic map distance between the centre of the window and the focal functional element, assuming either the pedigree-based recombination map for *M. musculus* constructed by Cox et al. (2009) or a recombination map estimated using linkage disequilibrium (LD) in the *M. m. castaneus* genome (Appendix). We excluded analysis windows that had a scaled genetic distance of *4N_e_r* < 1, because downstream analyses assume that sites are not tightly linked. All remaining analysis windows were collated into genetic distance bins. The average number of pairwise differences (i.e. nucleotide diversity) and nucleotide divergence from *Rattus rattus* were calculated for each genetic distance bin. For all analyses, we only examined the autosomes. Downstream analyses were sensitive to the assumption of a single mutation rate, so we excluded hypermutable CpG-prone sites from our analyses, identified as sites that were preceded by a C or succeeded by a G in the 3’ to 5’ direction in the reference genome (see Materials and Methods).

The choice of recombination map had a substantial effect on the profiles of average nucleotide diversity observed around protein-coding exons and CNEs (Figures S1, S2). When assuming the pedigree-based Cox map, we found that nucleotide diversity was slightly higher in the immediate flanks of both exons and CNEs (with distances calculated using the estimated recombination frequency), and lower in regions far from functional elements, compared to results with the LD-based map (Figure S1, S2). These differences are consistent with the possibility that the Cox map, which was constructed with a far smaller number of markers than the LD-based map, does not fully capture genomic regions that have either unusually low or high recombination rates. A possible consequence of this would be that analysis windows at intermediate distances from functional elements may appear to be more tightly linked to those elements than they actually are. This is supported by differences in numbers of sites falling into various genetic distance bins between the pedigree-based and LD-based recombination maps (Figures S1 and S2). However, both selective sweeps and BGS can induce LD, and may thus downwardly bias recombination rate estimates obtained using LD-based approaches (Clark et al., 2010). For this reason, we focus on results obtained assuming the pedigree-based Cox map for the remainder of the paper. We present parallel analyses, which assume the LD-based map, in the supplement and describe differences between the respective conclusions in the Discussion.

All mouse taxa exhibited dips in nucleotide diversity around protein-coding exons and CNEs (Figure 1). To quantify the relative reduction in diversity for each taxa, we calculated *π/π_Ref_*, the ratio of π to the average *π* at distances greater than *4N_e_r* = 1,500 and less than *4N_e_r* = 2,500 for exons, and distances greater than *4N_e_r* = 150 and less than *4N_e_r* = 250 for CNEs. The distances for determining *π_Ref_* were chosen based on where *π* began to flatten off with increasing distance from functional elements. Despite the existence of large differences in genome-wide diversity between the taxa, troughs in *π/π_Ref_* around exons and CNEs were very similar among mouse lineages (Figures 1, S1, S2). Nucleotide diversity was reduced by 20-30% and 10-20% around protein-coding exons and CNEs, respectively (Figure 1). The dips in diversity extended to genetic distances of up to approximately *4N_e_r* = 1,000 around exons, but only to *4N_e_r* = 100 around CNEs (Figure 1). Consistent with Halligan et al. (2013), we observed little reduction in between-species divergence around the edges of protein-coding exons, suggesting that mutation rate variation is not a substantial driver of the observed dips in diversity. However, in the immediate flanks of CNEs, we observed a trough in divergence. This may be explained if the *phastCons* approach used to identify CNEs did not readily identify weakly conserved sequences at the edge of more strongly conserved blocks. This would imply that that some sites subject to purifying selection tightly linked to the CNEs may have remained unannotated in our analysis. However, the troughs in nucleotide divergence around CNEs were substantially narrower than the corresponding troughs in diversity. This implies that reduced mutation rates or constrained sites may account for part of the diversity drop around CNEs, but do not explain all of it (Figure S1, S2).

**Figure 1.**
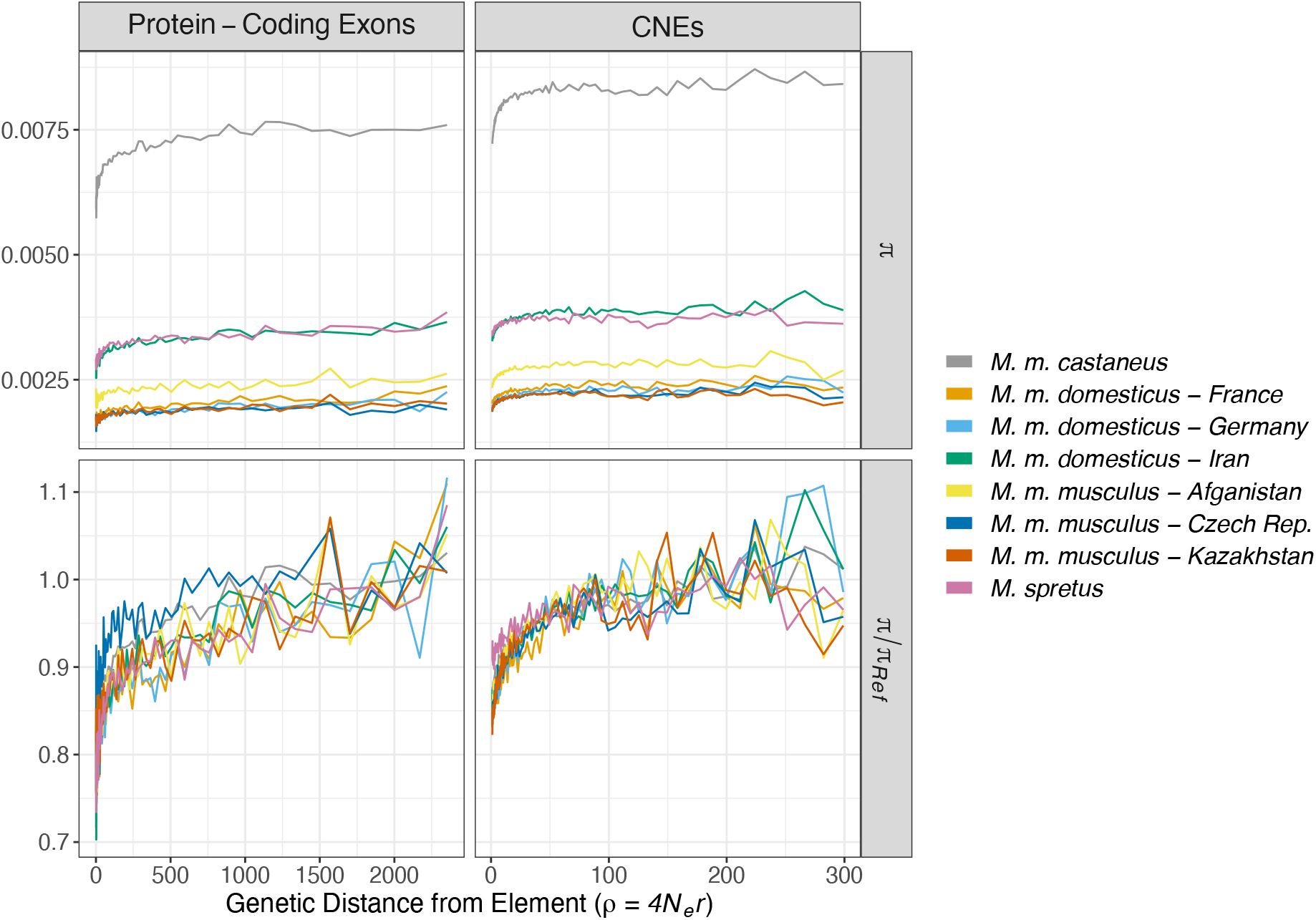
Nucleotide diversity (*π*) in regions surrounding protein-coding exons and CNEs in wild mice. Population-scaled recombination rates (*4N_e_r*) were calculated assuming the recombination map for *M. musculus* constructed by Cox et al. (2009). *π_Ref_* is the mean diversity calculated for sites far from functional elements.

An important caveat concerning the above analysis is that the mouse taxa thatare subject of the analysis are very closely related, i.e. it has been estimated that the *M. musculus* sub-species complex began to diverge around 350,000 years ago (Geraldes et al, 2011). Furthermore, Geraldes et al (2011) found extensive shared nucleotide variation among the sub-species, and that the average *F_ST_* among the members of the sub-species complex ranged from 0.43 to 0.72. Thus, patterns of polymorphism identified in the species are likely to be highly non-independent, and differences in *π* between the groups presumably reflect fluctuations in population sizes.

### Nucleotide polymorphism and divergence in wild mouse genomes

A first step for determining whether there was a consistent signal of natural selection across the mouse genomes, was to identify three classes of functional sites and two classes of putatively neutral sites as follows. For protein-coding gene orthologues between mouse and rat, we identified 0-fold degenerate nonsynonymous sites and UTRs, and used 4-fold degenerate sites as a neutral comparator. Protein-coding sites within the binding motifs of exonic splice enhancers including synonymous sites appear to be subject to purifying selection, implying that 4-fold sites located within them cannot be considered as neutral (Savisaar and Hurst 2018). We therefore excluded all synonymous and non-synonymous sites located within regions that matched such binding motifs. The total numbers of polymorphic and invariant sites that passed filtering for each of the mouse taxa are detailed in Supplementary File 1. We identified sites in the upstream and downstream flanks of individual CNEs for use as neutral comparators (see Methods).

To determine whether there was a consistent signal of natural selection, we assessed nucleotide diversity and lineage-specific divergence for the three classes of putatively functional sites (Figure 2). In all cases, functional site diversity and divergence were lower than for their putatively neutral counterparts, consistent with the action of purifying selection (Figure 2). Note that the *M. m. castaneus* data have been analysed in this way before (Halligan et al. 2013; Booker and Keightley 2018) and, as previously reported, *M. m. castaneus* had the highest nucleotide diversity of all *Mus* taxa surveyed (Figure 2; Harr et al. 2016). However, nucleotide divergence reported is the lineage specific divergence accumulated since the focal taxa began to diverge from *Mus famulus*, so that divergence estimates for the various mouse taxa are highly non-independent because of shared histories.

**Figure 2.**
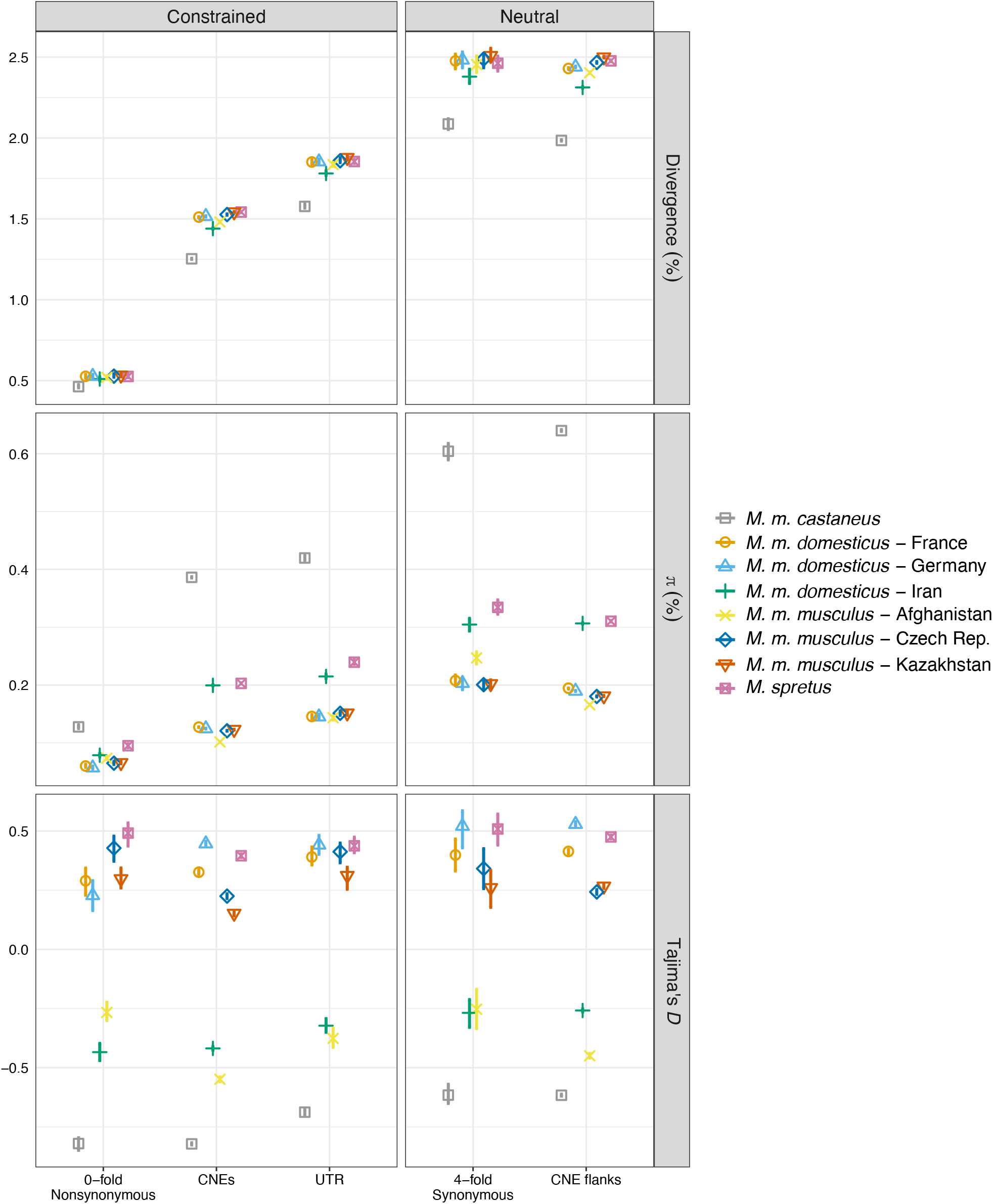
Population genetic summary statistics for three classes of putatively functional sites in the mouse genome and two putatively neutral comparators. Nucleotide diversity (*π*) and Tajima’s *D* are also shown. Error bars represent 95% confidence intervals based on 100 bootstrap samples. Those not visible are shorter than the height of the points.

All populations had nonzero Tajima’s *D* for putatively neutral sites, indicating the presence of either non-equilibrium population dynamics or genome-wide effects of selection (Figure 2). Mouse populations from Western Europe and Kazakhstan exhibited positive Tajima’s *D* for all classes of sites (Figure 2), consistent with a recent history of admixture between different populations or population bottlenecks (Charlesworth and Charlesworth 2010, pp.290-291). A population structure analysis of the mice analysed in this study did not suggest strong admixture between the sampled groups (Harr et al. 2016), but we cannot rule out the possibility of admixture with other unsampled mouse populations. *M. m. castaneus* and populations sampled in Iran and Afghanistan had strongly negative Tajima’s *D* values, consistent with recent population expansion or a genome-wide effect of recurrent selective sweeps (Charlesworth and Charlesworth 2010, pp.290, 414). Indeed, simulations modelling *D. melanogaster* populations have shown that recurrent, strong selective sweeps can induce negative Tajima’s *D* as large as −0.156 at synonymous sites (Campos and Charlesworth 2019). It worth noting that Tajima’s *D* is sensitive to the number of individuals and nucleotides analysed (Simonsen et al. 1995), which vary among the mouse taxa, so it is not straightforward to interpret differences in demographic history or strength of selection from these data.

### The distribution of fitness effects for deleterious mutations inferred from the uSFS

To parameterise a model of BGS, we estimated the distribution of fitness effects (DFE) for deleterious mutations in each of the mouse taxa by fitting a model of mutation-selection-drift balance to the unfolded site frequency spectrum (uSFS). The uSFS is a vector of 0, 1, 2, …, *k* counts of derived alleles, where *k* is the number of haploid genomes sampled. Estimates of the DFE can be obtained by contrasting the uSFS for a selected class of sites and a neutral comparator. Here, we estimated the uSFS for the three classes of functional sites and their putatively neutral comparator sequences. For each class of sites, we fitted a gamma distribution of deleterious mutational effects using *polyDFE* (v2; Tataru and Bataillon 2019). Tataru et al. (2017) showed that *polyDFE* provides robust estimates of the DFE for deleterious mutations based on the uSFS if a discrete class of beneficial mutations is also inferred. While the inferred beneficial mutation parameters are often spurious, including them seems to improve inference of the DFE for deleterious mutations (Booker 2020). Finally, while the gamma distribution is an arbitrary choice of model, and other probability distributions may give better fits to the data, it can capture the important features of the DFE, even if the underlying distribution is multi-modal (Kousathanas and Keightley 2013). Additionally, using the same probability distribution across taxa provides a consistent framework for comparing molecular evolution in the different mouse groups.

The estimated DFEs were all highly leptokurtic and had similar estimated parameters across the different taxa (Figure 3; Supplementary Table 2). Using *polyDFE*, the DFE is estimated in terms of the scaled selection coefficient for deleterious mutations, *2Nesd*, where *sd* is the reduction in fitness experienced by an individual homozygous for the mutation (which is assumed to be semi-dominant). Figure 3 shows the distribution of effects of deleterious mutational effects discretised into three ranges; nearly neutral mutations with 2*N_e_s_d_* < 1, mildly deleterious mutations with 1≤2*N_e_s_d_* < 10 and mutations with 2*N_e_s_d_* ≥ 10. Consistent with previous studies, amino-acid changing mutations were found to have the highest probability of having strongly deleterious effects (Halligan et al. 2013) and non-coding elements (UTRs and CNEs) had higher fractions of nearly neutral mutations (Figure 3). For 0-fold degenerate sites and CNEs, *M. m. castaneus* had the smallest proportion of nearly neutral variants among the taxa. The DFE inferred for the *M. m. musculus* sample from Afghanistan had the highest proportion of strongly deleterious mutations in UTRs, but this may reflect sampling error, since there were only 6 individuals and the population had among the lowest levels of nucleotide diversity (Figure 2).

**Figure 3.**
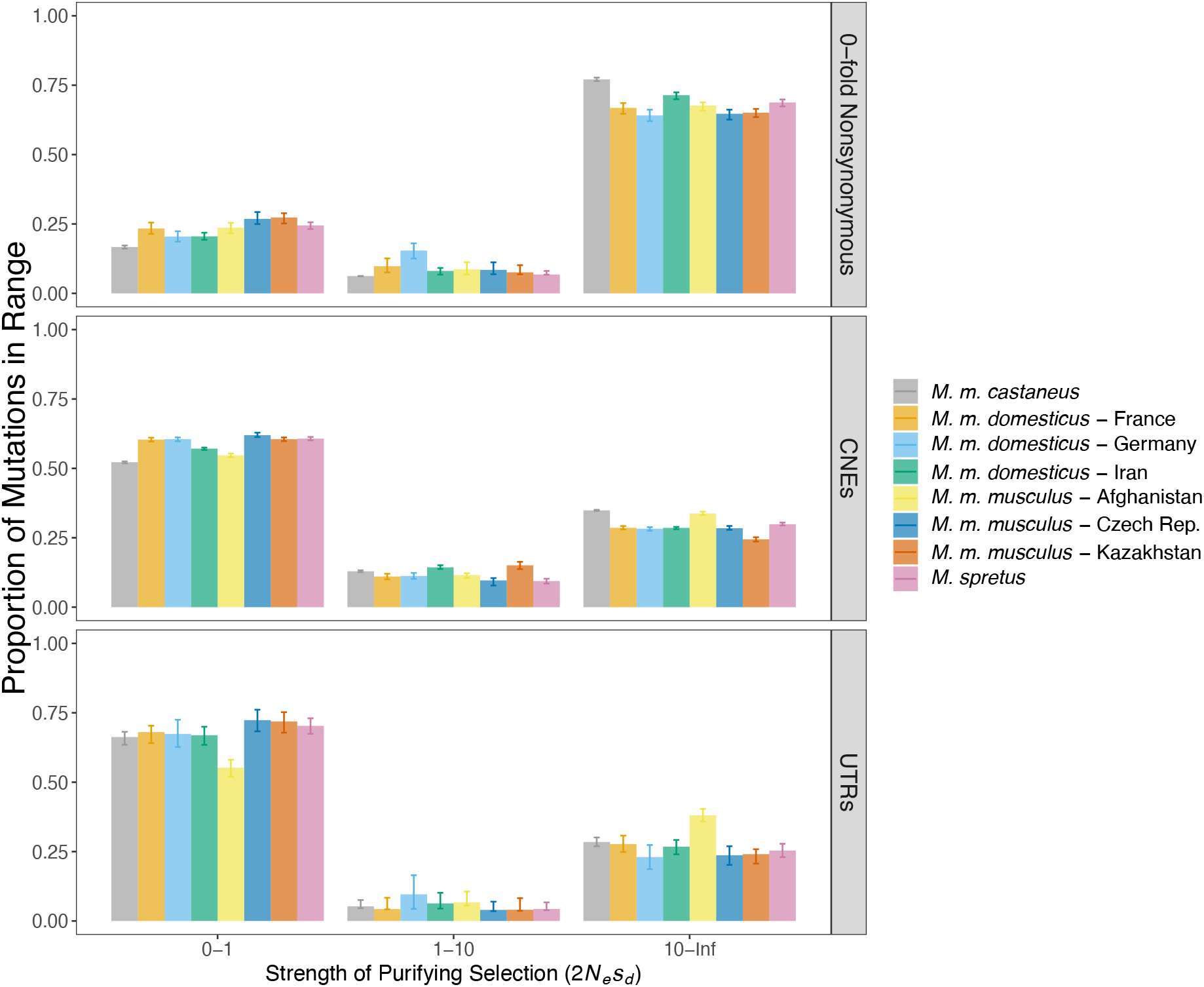
Graphical representation of the distribution of fitness effects of deleterious mutations for three classes of functional sites in wild mice. The figure shows the proportion of mutations falling into three ranges of effect size assuming a gamma DFE for each taxa and class of sites. Error bars indicate the 95% range based on 100 bootstrap replicates.

### The contribution of background selection to patterns of diversity around functional elements

Using the inferred DFE parameters for deleterious mutations, we can estimate the contribution of BGS to reductions in nucleotide diversity across the mouse genome. Specifically, we used simulations modelling *M. m. castaneus* to estimate the contribution of BGS to troughs in diversity observed around functional elements (Figure S3). Our simulations incorporated recombination rate variation (assuming either the pedigree-based or LD-based recombination maps, see below), the distribution of exons, UTRs and CNEs in the mouse genome, and the distributions of fitness effects for deleterious mutations estimated for those elements. We estimated values of *π* around both exons and CNEs from simulated data in the same manner as for the empirical data. Using the simulation results, we estimated the reduction in diversity caused by background selection, *B*, around functional elements for each genetic distance bin. We calculated *B* = *π/π*_0_, where *π* is the nucleotide diversity observed in the simulation and *π_0_* is the neutral expectation. As we found previously (Booker and Keightley 2018), BGS could not fully explain the reductions in diversity observed around protein-coding exons or CNEs (Figure S3).

Inferences about the strength of BGS made under the assumption of constant population size be misleading if there has been recent population size change. For example, a population bottleneck may lead to the accumulation of weakly deleterious mutations if drift overwhelms selection. As population size increases after a bottleneck, rapid purging of weakly deleterious mutations can occur, leading to deviations from the expectations of standard models of BGS, which assume constant population size (Torres et al. 2020; Johri et al. 2021). We have previously inferred a model of demographic history for *M. m. castaneus*, which suggested that population size has recently increased following a bottleneck (Booker and Keightley 2018). We performed an additional set of simulations incorporating this demographic history, but found that the relative reductions in diversity around both protein-coding exons and CNEs were very similar to those observed under constant population size (Figure S4). Note that the trajectory of the demographic history (bottleneck followed by recovery) we inferred may be an artefact of BGS (Ewing and Jensen 2016; Johri et al. 2021). However, we proceeded with our analysis assuming estimates of *B* for a constant population size, because the variations in *B* around exons and CNEs were very similar with or without population size change.

### Parameters of beneficial mutations obtained from patterns of nucleotide diversity

We estimated the parameters of beneficial mutations occurring in protein-coding and gene regulatory regions by fitting a model that combines the effects of BGS and recurrent selective sweeps to troughs in average nucleotide diversity around functional elements (see Materials and Methods). The model quantifies the reduction in neutral diversity surrounding the average exon or CNE, assuming that they are 150bp and 52bp long, respectively. A key parameter in the model is *π_0_* = *4N_e_μ*, the nucleotide diversity expected under neutrality in the absence of selection at linked sites, where *N_e_* is the effective population size and *μ* is the mutation rate per basepair. Estimation of *π_0_* is problematic, however, and *π_0_* may even be unobservable in empirical data, given the ubiquity of selection at linked sites (Kern and Hahn 2018). In the empirical data, *π* levelled off at different values for protein-coding exons and CNEs (Figure 1, S1, S2). However, our simulations predicted that *B* should plateau at around 0.95 in genomic regions surrounding both protein-coding exons and CNEs (Figure S3). *B* was not predicted to plateau at 1.0 in our simulations, because we modelled the distribution of all functional elements in the genome, so that a site may be influenced by BGS generated by many surrounding elements. Our simulations did not model sweeps, so simply dividing empirical *π* by our estimated *B* would give an underestimate of *π_0_*, because the reduction in diversity caused by positive selection was not included. When analysing variation in *π*, we therefore assumed values of *π_0_* = 0.0081 and 0.0091 for protein-coding exons and CNEs, respectively, to reflect the different levels at which diversity plateaued.

We proceeded to fit models combining BGS and selective sweeps to the troughs in diversity around protein-coding exons and CNEs in *M. m. castaneus* assuming various models for the effects of advantageous mutations. We estimated the strength of selection acting on new, semi-dominant beneficial mutations as *γ_a_* = 4*N_e_s_a_*, where *s_a_* is the increase in relative fitness experienced by heterozygotes. We also estimated *p_a_*, the proportion of new advantageous mutations in a functional element. We found that a model with two classes of advantageous mutations gave a better fit than a single class of mutations or an exponential distribution of effects (as judged by AIC; Supplementary File 3). This result held regardless of the recombination map that was assumed (Supplementary File 3).

For both protein-coding exons and CNEs, we found that the best fitting model included a class of strongly advantageous mutations and a class of more mildly beneficial mutations (Table 1). When assuming the pedigree-based Cox map, we estimated the scaled fitness effects of the strongly selected class (*γ_a_*) to be 6,200 and 1,900 for protein-coding exons and CNEs, respectively. The proportions of mutations with these selection coefficients were 9 x 10^-6^ and 3 x 10^-4^, respectively. The more mildly beneficial class of mutations inferred for protein-coding exons and CNEs had scaled effects of 210 and 7.0, respectively, and the proportion of mutations with these effects were 3.5 x 10^-4^ and 1.8 x 10^-2^, respectively. In the case of CNEs, although two classes of advantageous mutational effects gave the best fit to the data, the coefficient of variation for the parameter estimates of the mildly selected class was large, and evidence for mildly beneficial mutations is fairly weak in this case (Table 1).

**Table 1.** Parameters of positive selection in *M. m. castaneus* estimated by fitting a model of selective sweeps and background selection to troughs in diversity around functional elements. The frequency (*p_a_*) and scaled selection coefficients (*γ_a_*) for the two classes of advantageous effects are given. Standard errors are shown in square brackets below point estimates.

| Element | $\gamma_{a,1}$ | $p_{a,1}$ | $\gamma_{a,2}$ | $p_{a,2}$ |
| --- | --- | --- | --- | --- |
| Protein-Coding Exons | 6,170<br>[2,650] | $0.80 \times 10^{-5}$<br>[$0.50 \times 10^{-5}$] | 208<br>[105] | $3.50 \times 10^{-4}$<br>[$1.80 \times 10^{-4}$] |
| CNEs | 1,910<br>[673] | $1.30 \times 10^{-5}$<br>[$0.60 \times 10^{-5}$] | 7.00<br>[3.50] | $1.78 \times 10^{-2}$<br>[$1.29 \times 10^{-2}$] |

The choice of recombination map strongly affected the estimated selection parameters obtained. Use of the pedigree-based Cox map resulted in estimated selection coefficients that were typically smaller than those obtained when assuming the LD-based recombination map (Supplementary Table 3). This is because we found the troughs in diversity around both exons and CNEs were shallower when calculating genetic distances using the pedigree-based map than when using the LD-based map (Figure S1, S2).

BGS appears to contribute to the troughs in diversity around both protein-coding exons and CNEs and causes an overall reduction in neutral diversity (Figure 4). Ignoring the contribution of BGS (i.e. by setting *B* to 1.0 when fitting Equation 4 to the diversity troughs) resulted in a much poorer model fit (Supplementary File 3). In the absence of BGS, the selection coefficients for advantageous mutations required to explain the observed data are, as expected, far higher (Supplementary File 3).

**Figure 4.**
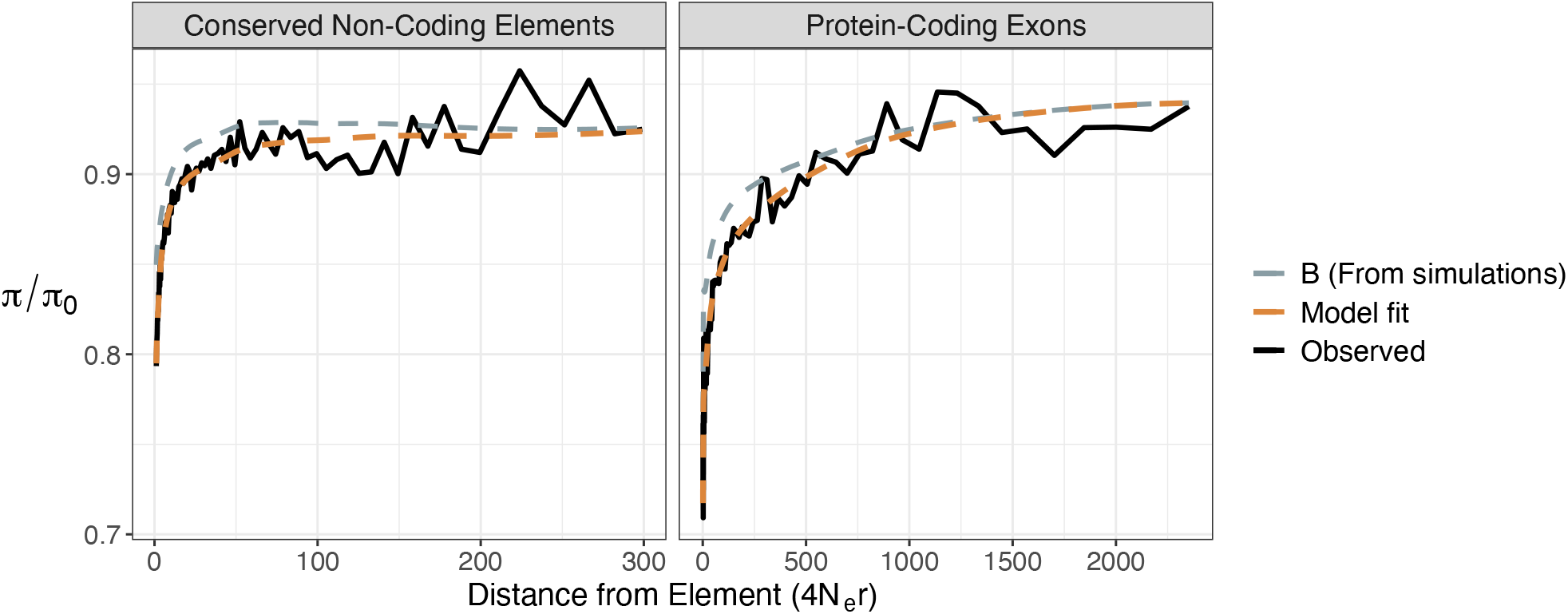
The reduction of scaled nucleotide diversity around protein-coding exons and CNEs in *M. m. castaneus*, predicted by fitting a model combining the effects of background selection and selective sweeps to the observed data. Genetic distances were calculated assuming the pedigree-based recombination map constructed by Cox et al. (2009). The effect of background selection (*B*) was estimated using simulations.

We did not include gene conversion events in our analysis, because gene conversion tracts, which have an estimated mean length in mice of 135bp (Paigen et al. 2008), are relatively short compared to the genetic distances we analysed (up to 100,000bp and 5,000bp for exons and CNEs, respectively). Furthermore, the ratio of the rates of gene conversion and crossover events has been estimated to be 0.105 in mice (Paigen et al. 2008). Overall, gene conversion is expected to contribute little to the net frequency of recombination between neutral and selected sites.

### The relative contribution of adaptive substitutions in protein-coding and regulatory regions to fitness change in mice

An important goal of evolutionary biology is to understand the extent to which protein-coding and regulatory elements contribute to phenotypic evolution (King and Wilson 1975; Wray 2007; Stern and Orgogozo 2008; but see Hoekstra and Coyne 2007). Using our estimated selection parameters, we can parameterise the following model of the rate of fitness change per generation (*ΔW*) brought about by the fixation of advantageous mutations. For a particular class of sites, assume there are *η_a_* nucleotides in the genome at which new mutations occur at rate *μ* per nucleotide site per generation. If the size of the breeding population is *N*, then 2*Nμ* new mutations enter the population each generation. We assume that a proportion of the new mutations, *p_a_*, is strongly advantageous, with a selection coefficient of *s_a_* in heterozygous carriers. When the effectiveness of selection exceeds that of genetic drift (*2N_e_s_a_* > 1), the fixation probability is approximately 2*s_a_* (Haldane 1927). Once fixed, advantageous mutations increase population mean fitness by *s_a_ /h*, where *h* is the dominance coefficient, giving the following expression:

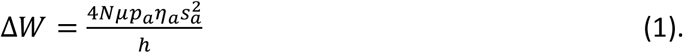

Since we are interested in the relative contribution to fitness change, and assumed that the average point mutation rate is the same for CNEs and protein-coding exons, we can thus ignore *μ* in Equation 1. Note that the above model is conceptually similar to an approach taken by Lynch et al. (1993) to model fitness change under mutational meltdown. We parametrized Equation 1 using our estimated selection parameters. Note that we estimated two classes of beneficial mutational effects for the two classes of functional elements. When parameterizing Equation 1, we summed the fitness contributions over the two classes of fitness effect inferred for each element. We calculated the ratio of *ΔW* for protein-coding exons and CNEs (*ΔW_Exons_/ ΔW_CNEs_*) as a measure of the relative contributions of the two types of elements to adaptive evolution (which also implicitly assumes the same *h* for all classes of mutation).

Our point estimates suggest that *ΔW* is larger for protein-coding regions than regulatory regions. However, it is notable that the total genomic rate of fixation of beneficial mutations is higher for CNEs than for coding regions (see also Halligan et al. 2013), but this reflects the fact that there are approximately three times as many CNE bases as non-synonymous bases in the mouse genome. Although the estimated genomic rate of fixation of beneficial mutations in CNEs is greater than that of protein-coding exons (Table 2), the average strength of selection acting on a new advantageous nonsynonymous mutation far exceeds that of CNEs (Table 2). Fitness change is proportional to the square of the effect size, so that the change in population mean fitness brought about by the fixation of advantageous mutations is substantially higher for protein-coding exons than for CNEs. This result is sensitive to the choice of recombination map, since we inferred stronger selection when assuming the LD-based map (Supplementary Table 3). Using a parametric bootstrap approach, we found that *ΔW_Exons_/ ΔW_CNEs_* was significantly greater than 1 when using the LD-based map, but not when assuming the pedigree-based map of Cox et al. (2009).

**Table 2.** Estimates of the change in fitness brought about by the fixation of advantageous mutations. Estimates were obtained assuming an effective populations size for *M. m. castaneus* of 420,000 and the selection parameters shown in Table 1.

| Recombination<br>Map | Element | $\Delta W_{Exons}/$<br>$\Delta W_{CNEs}$ | 95% Bootstrap<br>interval |
| --- | --- | --- | --- |
| LD-based<br>(*castaneus* map) | Protein-Coding Exons<br>CNEs | 23.11 | 11.08 - 54.67 |
| Pedigree-based<br>(Cox et al. 2009) | Protein-Coding Exons<br>CNEs | 2.94 | 0.211 - 46.36 |

### Selective sweeps and background selection in the mouse genome

The profiles of nucleotide diversity indicate the existence of pervasive effects of selection on diversity across the genome (Figure 1, Figure 2). By fitting a model of sweeps to the troughs in diversity around protein-coding exons and CNEs, while assuming that the troughs are partly caused by BGS, we estimated the parameters of positively selected mutations occurring in the two classes of element. Our analysis suggests that regulatory sequences experience a higher genomic rate of newly arising advantageous mutations than protein-coding sites. However, the trough in diversity around exons is both deeper and wider than what is observed around CNEs, and, accordingly, we found that protein-coding regions experience more strongly selected mutations than regulatory sequences. Using a different approach, Campos et al. (2017) came to a similar conclusion for *D. melanogaster* by comparing UTRs with the coding sequences of genes.

Due to non-independence among the various *M. musculus* sub-species, we only estimated parameters of positive selection for *M. m. castaneus*, the sub-species with the highest levels of diversity. Our selection parameter estimates for *M. m. castaneus* are fairly similar to estimates obtained for European *M. m. domesticus* in an earlier study (Teschke et al. 2008).

### Limitations and next steps

There are a number of caveats concerning our estimates of positive selection parameters. Firstly, we found that an exponential distribution of beneficial mutational effects provided a poorer fit to the troughs in diversity compared to a model with two discrete classes of effects (Supplementary File 3). However, the true DFE for advantageous mutations is almost certainly more complex than the simple models assumed. The approach used in this study was based on average nucleotide diversity across many sites, and we presumably had little power to infer a more complex model of the DFE for advantageous mutations. Secondly, we have assumed that all elements of a particular class share a common set of selection parameters. This is problematic, since CNEs could be composed of several categories, such as promoters and enhancers, which may be subject to different selective pressures. Indeed, different categories of protein-coding genes may also be subject to different selection pressures. For example, immunity genes in *D. melanogaster*, virus interacting proteins in humans and highly expressed genes in *Capsella grandiflora* appear to have higher rates of adaptive substitutions than the respective genome wide averages (Enard et al. 2014; Obbard et al. 2009; Williamson et al. 2014). Thirdly, for a single class of advantageous mutational effects, under the assumption that there is no interference among sweeps, the predicted reductions in diversity caused by selective sweeps can be modelled as a simple hyperbolic function (Equation 4). However, if the rate of sweeps is sufficiently high, and the rate of recombination is sufficiently low, selective interference can cause the rate of sweeps to be lower than predicted by a given strength of selection (Campos and Charlesworth 2019). This implies that the strength of positive selection would be overestimated by our methods. A bias in the opposite direction, which is likely to be more important for genomic regions with normal levels of recombination, is caused by deviations from one of the assumptions underlying Equation 4, i.e. that there is full recovery of nucleotide diversity between selective sweeps (Campos and Charlesworth 2019; Charlesworth 2020). This would lead to the effects of sweeps to be underpredicted. Incorporating more sophisticated models of selective sweeps into the inference framework is a logical next step.

The architecture of functional elements in the mammalian genome is such that a single exon or CNE is rarely far away from another functional element. When estimating the effects of sweeps on neutral diversity, we excluded all putatively functional sites from our analysis windows, but multiple linked elements may affect observed diversity at a given locus. For this reason, we did not estimate the strength of positive selection acting on UTRs, although sweeps in these elements are also likely to contribute to heterogeneity in *π* across the mouse genome, as has been found in *D. melanogaster* (Campos et al. 2017). The model fitted to the troughs in diversity assumes that selection is generated by a single, idealised exon or CNE. However, there is variation in the length of exons and CNEs across the genome. An analysis that models genome-wide heterogeneity in diversity while taking into account the locations of individual functional elements, similar to the method developed by Elyashiv et al. (2016) for *D. melanogaster*, could be a more powerful approach. Note that the approach of Elyashiv et al. (2016) might not be applicable in all situations, because it conditions the effects of sweeps on the locations of recent substitutions. In mice and humans, patterns of diversity around nonsynonymous substitutions are indistinguishable from the patterns of diversity around synonymous substitutions (Hernandez et al. 2011; Halligan et al. 2013). Developing a chromosome-wide analysis that conditions the effects of sweeps on the locations of genomic elements rather than substitutions may be a useful avenue for further research.

The model of sweeps we assumed involves positive selection acting on *de novo* mutations – the so-called ‘hard’, or ‘classic’ sweep model. Studies in humans and *Drosophila* have, however, suggested that ‘soft’ sweeps are common (Garud et al. 2015; Garud and Petrov 2016; Schrider and Kern 2016; but see Harris et al. 2018). Soft selective sweeps occur when advantageous alleles present in multiple copies in the population spread to fixation, which can occur if selection acts on standing genetic variation or if multiple copies of the selected allele arise independently (Hermisson and Pennings 2017). Additionally, adaptation acting on quantitative traits subject to stabilising selection may generate partial sweeps, because changes in allele frequencies at many loci can rapidly alter mean phenotypes, without necessarily causing fixations (Pritchard et al. 2010; Jain and Stephan 2017). The profiles of the reductions in diversity around soft and partial sweeps differ from those expected under hard sweeps, and if either of the alternative types of sweep were common, the assumption of a hard sweep model could result in spurious parameter estimates (Elyashiv et al. 2016). Finally, the trough in diversity around a selective sweep in a structured population is expected to be shallower than in a panmictic population because of the longer time taken to reach fixation (Barton 2000; Santiago and Caballero 2005). If beneficial alleles are frequently introduced via migration, we may therefore underestimate the strength of selection.

Finally, it is important to note that CNEs are generally expected to represent regulatory sequences that are deeply conserved. It has been demonstrated that the evolution of regulatory elements is more dynamic than that of coding sequences, with major gains of new regulatory elements having occurred in vertebrate and mammalian evolution (Mikkelsen et al. 2007; Lowe et al. 2011). If more recently acquired regulatory elements, which may be absent from the CNE dataset, experience stronger or more frequent adaptive substitutions, it is possible that we have underestimated the contribution of regulatory changes to adaptive evolution. For instance, a recent gain of a new regulatory element might have been caused by relatively strong positive selection acting on the element as a whole, resulting in a single sweep event. This would fall outside the inference framework developed here.

It seems likely that adaptation does not fit any one particular mode, but rather different functional elements will be subject to a mixture of different types of sweep that may vary depending on the genomic region. For example, adaptation may more commonly act on standing variation in regulatory regions simply because they harbour greater nucleotide diversity than nonsynonymous sites (Figure 2).

## Conclusions

In this study, we have shown that multiple wild mouse taxa exhibit patterns of genetic diversity and divergence that are consistent with the action of natural selection. Furthermore, we have shown that strong positive selection can explain the dips in diversity around protein-coding exons and CNEs in *M. m. castaneus*. Finally, even though the framework we have adopted here is incapable of distinguishing different modes of positive selection such as adaptation, sexual selection and various forms of competition, the estimated parameters of positive selection suggest that mutations in protein-coding regions may contribute more to the rate of change in fitness under positive selection than regulatory mutations.

## Materials and Methods

### Genomic data

We re-analysed previously published genome sequences for the 54 wild-caught *Mus musculus* individuals described in Harr et al. (2016) and the 10 *M. m. castaneus* individuals and the *M. famulus* individual originally described in Halligan et al. (2010, 2013). The mouse samples belonged to three species: *Mus spretus, Mus musculus* and *Mus famulus*. The *M. spretus* individuals (*n* = 8) were from Madrid, Spain. The *M. musculus* individuals are composed of samples from the sub-species *M. m. domesticus, M. m. musculus* and *M. m. castaneus*. Three populations of *M. m. domesticus* were sampled (Massif Central, France, *n* = 8; Cologne-Bonn, Germany, *n* = 8; Ahvaz, Iran, *n* = 8) and three populations of *M. m. musculus* were sampled (Afghanistan, *n* = 6; Studenec, Czech Republic, *n* = 8; Mazar-e-Sharif, Kazakhstan, *n* = 8). We also analysed 10 *M. m. castaneus* described by Halligan et al. (2010, 2013), sampled in Himachal Pradesh, India. The one *M. famulus* individual, originated in Southern India, though Halligan et al. (2013) obtained it from the Montpellier Wild Mice Repository.

Harr et al. (2016) published and made available the variant calls obtained from the *M. musculus* samples described above in the form of VCF files. However, Harr et al. (2016) did not include invariant sites in their VCFs; for our purposes we required this information, so we re-called variants from their processed BAM files, available at http://wwwuser.gwdg.de/~evolbio/evolgen/wildmouse/. The data had been processed according to the GATK version 3 best practices pipeline, up to the step prior to variant calling. Briefly, all sequencing reads had been mapped to the *mm10* genome using *bwa-mem* (Li 2013). Reads were then sorted, merged and PCR duplicates were marked using *picardtools* (https://broadinstitute.github.io/picard/). Base Quality Score Recalibration was then applied using the dbSNP resource for mice (https://www.ncbi.nlm.nih.gov/snp) to produce analysis-ready alignments in BAM format. We generated BAM files for the *M. m castaneus* data and the *M. famulus* mice using the same procedure using FASTQ files downloaded from the European Nucleotide Archive (accession number PRJEB2176). For each of the mice, we called variants separately using the HaplotypeCaller tool from GATK3.7 (McKenna et al. 2010), with the options “–*emitRefConfidence BP_RESOLUTION –max-alternate-alleles* 2”, and made population-specific VCF files using the GATK tools *combineGVCFs* and *genotypeGVCFs*. We restricted all analyses to autosomal sites.

### Outgroup information and CpG sites

In this study we used *M. famulus, Mus pahari* and *Rattus norvegicus* as the outgroup species. For each of the mouse taxa described above and each outgroup, we created a synthetic mm10-length reference genome by replacing mm10 alleles with the major allele of the variant call set. In addition, we constructed a synthetic genome for *R. norvegicus* by replacing mm10 alleles with the homologous positions in the rat genome using the UCSC reciprocal best alignments between rn6 and mm10 (available at: ftp://hgdownload.cse.ucsc.edu/goldenPath/rn6/vsMm10/reciprocalBest/) using custom Python scripts. For an additional outgroup, more closely related to *Mus musculus* than the rat, we obtained the homologous alleles from *Mus pahari* at mm10 positions using the ENSEMBL pairwise alignments between the *M. pahari* reference sequence (Thybert et al. 2018) and mm10 (available at: ftp://ftp.ensembl.org/pub/release-90/maf/ensembl-compara/pairwise_alignments/).

CpG sites have higher rates of spontaneous mutation than non-CpG sites, and identifying and excluding CpG-prone sites is a conservative way of reducing the impact of CpG hypermutability on analysis of population genomic data (Gaffney and Keightley 2008). For each of the rodent taxa, we used the synthetic mm10-length reference genomes to identify the locations of CpG-prone sites, defined as those sites in our synthetic references that were preceded by a C or followed by a G in the 5’ or 3’ direction, respectively. All analyses presented in this paper excluded CpG-prone sites.

### Annotations and identifying conserved non-coding elements

We downloaded the list of mouse-rat orthologs from https://www.ensembl.org/biomart/ and extracted the annotations for each from version 38.93 of ENSEMBL (Mus_musculus.GRCm38.93.gtf.gz; Howe et al. 2021). For each of the orthologs, we identified the positions of 0-fold degenerate nonsynonymous and 4-fold degenerate synonymous sites using the synthetic genomes for each of the mouse taxa and the outgroups described above. The locations of 5’ and 3’ untranslated regions (UTRs) were retained for downstream analyses. We also retained a list of all exonic positions in the mouse genome, regardless of orthology, for the purposes of filtering out functionally constrained sites in downstream analyses.

There is evidence that synonymous sites within exonic splice enhancers (ESEs) in humans are subject to purifying selection, and that ignoring ESEs can bias analyses that rely on the assumption that synonymous sites evolve neutrally (Savisaar and Hurst 2018). Savisaar and Hurst (2018) identified putative ESEs by comparing human gene sequences against various lists of ESE motifs. They found that synonymous sites in regions matching ESE motifs had lower nucleotide diversity than those outside of putative ESEs. We identified the locations of potential ESEs in protein-coding genes orthologous between mice and rat using the merged list of ESEs described in Savisaar and Hurst (2018) (kindly provided by Rosina Savisaar). For each of the mouse-rat orthologs, we extracted the gene sequence and performed a string search against the list of ESE motifs. We recorded the genomic position of each region matching an ESE motif and used them to filter out the affected coding sites in downstream analysis.

We identified conserved non-coding elements (CNEs) in murid rodents using a 40-way alignment of placental mammals downloaded from UCSC (http://hgdownload.cse.ucsc.edu/goldenPath/mm10/multiz60way/). To avoid ascertainment bias, the mouse and rat genomes in the 40-way alignment were converted to the character “N” prior to calling conserved elements, following Williamson et al. (2014). We ran *phastCons* with the following arguments --expected-length=45 --target-coverage=0.3 --rho=0.31. To identify CNEs, we masked all exonic regions from the resulting file of *phastCons* elements using the complete list of annotations from the 38.93 database (see above). The scripts and full pipeline used to identify CNEs are available at https://github.com/rorycraig337/mouse_mm10_conserved_elements.

For each CNE identified in this way, we obtained the location of their flanking sequences, which we used as neutral comparators in downstream analysis. For each CNE, we recorded the locations of two loci of equal length upstream and downstream of the focal element, offset by 500bp. We merged overlapping CNE-flanks and masked out sites that overlapped any CNE or exonic sites.

We analysed the mouse genomes assuming the pedigree-based genetic map of *Mus musculus* constructed by Cox et al. (2009). The Cox map was constructed using data from 3,546 meioses observed in crosses of common laboratory strains. The markers genotyped by Cox et al. (2009) were mapped to the mm9 reference genome, but in the present study we converted the mm9 coordinates to mm10 positions as follows. The Cox map was downloaded from the Jackson

Laboratory website (http://cgd.jax.org/mousemapconverter/). The SNP positions of the Cox map were then extracted and converted to mm10 positions using the online UCSC LiftOver tool (https://genome.ucsc.edu/cgi-bin/hgLiftOver). The physical distances between the mm10 SNP positions were then converted to units of genetic distance using the Jackson Laboratory’s conversion tool (http://cgd.jax.org/mousemapconverter/). We also analysed the mouse genomes using an LD-based recombination map inferred from the sample of *M. m. castaneus* individuals, as described in the Appendix.

### Mouse analysis – Patterns of nucleotide diversity around selected sites

For each of the *M. musculus* sub-species and *Mus spretus*, we examined patterns of nucleotide diversity around protein-coding exons and CNEs. From the edges of protein-coding exons (CNEs), polymorphism data and divergence from the rn6 rat reference genome were extracted in windows of 1Kbp (100bp) extending to distances of 100Kbp (5Kbp). Analysis windows only extended to the midway point between adjacent elements. Sites within the exons of protein-coding genes or CNEs were excluded from analysis windows. The genetic distance between an analysis window and a focal element was calculated either from the pedigree-based genetic map constructed using common lab strains of *M. musculus* (Cox et al. 2009) or the linkage disequilibrium (LD) based recombination map for *M. m. castaneus*. The SFS and divergence from were recorded for each analysis window. Analysis windows were then binned based on the genetic distance from the focal element, and the SFS and divergence from individual windows were collated. Because LD-based and pedigree-based recombination maps have different features and shortcomings (see Results), we performed analyses based on both genetic maps.

### Estimating the unfolded site frequency spectrum, summary statistics and the distribution of fitness effects

We analysed genetic variation for five different classes of sites in the genome, i.e. the 0-fold and 4-fold degenerate sites and UTRs of protein-coding genes, CNEs and CNE-flanks. For each class of sites, we inferred the unfolded site frequency spectrum (uSFS), which is the distribution of derived allele frequencies in the mouse samples. The uSFS was inferred by maximum likelihood using the two-outgroup method of Keightley and Jackson (2018) using *Mus famulus* and *Mus pahari* as outgroups. We compared the fit of 1-, 2- and 6-parameter mutation rate models using *est-sfs* (v2.03; Keightley and Jackson 2018). Consistently, a model with 6 mutation rate parameters (i.e. the R6 model from Keightley and Jackson 2018) provided the best fit to the data (as judged by model likelihoods), but the uSFS and lineage specific divergences that were estimated under the 2-parameter and 6-parameter models were almost identical in all cases (Supplementary File 1), so we have used the results from the 2-parameter model in our analyses for all taxa. For each taxon and class of sites, we performed 100 bootstraps, sampling genes or CNEs with replacement. We inferred the uSFS for each bootstrapped dataset as above. For each class of sites, we calculated nucleotide diversity (*π*) and Tajima’s *D* from the inferred uSFS for each bootstrap sample.

Estimates of the distribution of fitness effects (DFE) were obtained by analysing the unfolded site frequency spectrum for each class of functional site using *polyDFE* v2 (Tataru and Bataillon 2019). For 0-fold sites and UTRs, we used 4-fold degenerate synonymous sites as the neutral comparator, and for CNEs we used CNE-flanks. Using *polyDFE2*, we fitted a gamma distribution of deleterious mutations effects and a single class of beneficial mutations (using the -model B option). We excluded between species divergence from the analysis using the “-w” option. We fitted the uSFS data for each of the bootstrap replicates described above.

### Simulating background selection

There is substantial evidence that background selection (BGS) contributes to troughs in diversity around protein-coding exons and CNEs (Halligan et al. 2013; Booker and Keightley 2018). For our analysis, we therefore required estimates of the effect of BGS on neutral diversity, *B*, at varying distances from functional elements. Estimates of *B* were included as covariates when fitting a model of selection at linked sites. However, when purifying selection is weak (*γd* < 5) analytical formulae for calculating *B* over-predict the effects of BGS (Gordo et al. 2002; Good et al. 2014), and weakly deleterious mutations appear to comprise a large fraction of the DFEs for mice (Figure 3 and Halligan et al. 2013). We therefore opted to obtain estimates of the variation in *B* from forward-in-time simulations that modelled the entire range of fitness effects inferred for mice.

We used *SLiM* v3.2 (Haller and Messer 2019) for this purpose. Following Booker and Keightley (2018), we incorporated the actual distribution of functional elements (the coding exons and UTRs of protein-coding genes, and CNEs) and the estimated recombination rates. 1 Mbp regions of the mouse genome were randomly sampled and the functional annotations in the sampled regions were used as the basis of a simulation replicate. The parameters of the gamma distributions of fitness effects for deleterious mutations estimated for 0-fold sites, UTRs and CNEs were used in the simulations for the respective elements. The recombination rate variation present in the sampled region of the mouse genome was included in the simulations using either the pedigree-based map from Cox et al. (2009) or the LD-based recombination map for *M. m. castaneus*. When assuming the Cox map, the recombination rates (in units of cM/Mbp) were scaled in the simulations by a factor of 420*r*, assuming *N_e_* = 420,000 for wild *M. m. castaneus*. In the case of the LD-based estimates of the recombination rate, population-scaled recombination rates (in units of *4N_e_r*) were simply divided by 4*N*, where *N* was the simulated population size. Populations of *N* = 1,000 diploid individuals were simulated for 20,000 generations. We set the mutation rate such that the neutral expectation *π_0_* = *4N_e_μ* = 0.01, based on the upper estimate of nucleotide diversity observed in the *M. m. castaneus* genome (Figure 1). Given the simulated population size of 1,000 diploids, *4N_e_μ* = 0.01 corresponded to a point mutation rate of *μ* = 2.5 x 10^-6^. We used the tree-sequence recording option in *SLiM* to record the genealogies of the simulated populations, so modelling neutral mutations in *SLiM* was not required. Instead, neutral mutations were added to the recorded coalescent trees at a rate *μ* using *PySLiM* (https://pyslim.readthedocs.io/en/latest/introduction.html). We sampled 200 haploid chromosomes from the population and extracted *B* = *π/π*_0_ as a function of genetic distance from both protein-coding exons and CNEs. Data were extracted from the simulated populations in the same way as for the empirical data. To obtain smoothed *B* values, we fitted LOESS curves to the average *π* observed around functional elements in the simulated data. We fitted LOESS curves using a span parameter of 0.3 and the number of sites contributing to each analysis bin as weights in R (v3.4.2).

### Model of recurrent selective sweeps and background selection

Our analysis is a modification of that of Elyashiv et al. (2016) and Campos et al. (2017), where expressions were described for the neutral diversity expected under the combined effects of BGS and sweeps. Consider a haplotype with a set of neutral sites linked to a site that is the target of positive selection. A new, semi-dominant advantageous mutation with heterozygous selection coefficient *s_a_* occurs at site *i* on the haplotype and spreads to fixation. Recombination between sites *i* and *k* uncouples the neutral and selected sites at rate *r_i,k_* per generation. The expected change in neutral diversity at site *k* (Δ*π_k_*), relative to its expectation in the absence of selection (*π_0_*) is given by

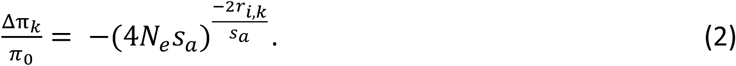

See Barton (2000), Charlesworth and Charlesworth (2010, p411) or Campos and Charlesworth (2019) for derivations of Equation 2. This approximation assumes that selection pressure on the advantageous allele satisfies *N_e_s_a_* >> 1, so that the sweep can be treated deterministically following an initial stochastic establishment phase. Under this assumption, the quantity −Δ*π_k_* in Equation 1 can be equated to the probability of a sweep-induced coalescent event at site *k* (Wiehe and Stephan 1993). For a particular class of functional elements (e.g. protein-coding exons), sweeps occur at a rate of *V_a_* = 2*μp_a_γ_a_* per nucleotide per generation (Kimura and Ohta 1971), where *μ* is the mutation rate per nucleotide site, *p_a_* is the proportion of new mutations that are advantageous and *γ_a_* is the scaled selection coefficient (4*N_e_s_a_*) for these mutations. If *V_a_* is sufficiently low, such that sweeps do not interfere with each other, the total probability of sweep-induced coalescence for a neutral site caused by selection at a linked functional element is:

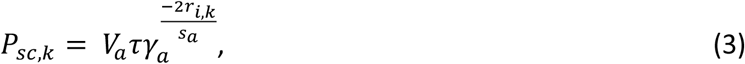

where *τ* is the number of sites in a particular class of functional element. In our analysis of data from wild mice, *r_i,k_* was measured from the end of a functional element to the centre of an analysis window.

The effects of background selection at site *k* can be represented by multiplying the effective population size by a factor *B_k_*. The probability of coalescence for a neutral allele affected by BGS is thus 1/(2*B_k_N_e_*). We assume that coalescent events caused by BGS and sweeps follow independent exponential distributions, so thar the rate of coalescence induced by the two processes is the sum of 1/(2*B_k_N_e_*) and *P_sc,k_*. We also multiply the sweep effect *P_sc,k_* by *B_k_* to reflect the reduction in the fixation probability of a new advantageous mutation as a result of the reduction in *N_e_* caused by BGS, following Kim and Stephan (2000). Simulations show that this may overestimate the effect of BGS on fixation probabilities (Campos and Charlesworth 2019), we thus compared selection parameters with and without including background selection.

Writing the reciprocal of the rate of coalescence at the neutral site under the combined effects of BGS and sweeps as *T_k_* (which is equivalent to the expected time to coalescence of a pair of alleles), and expressing it relative to the expected time to coalescence under neutrality (*T_0_*), we have:

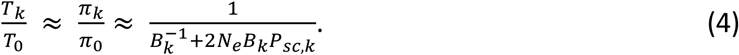

We estimated parameters of advantageous mutations by fitting Equation 4 to the relationship between nucleotide diversity and genetic distance from functional elements, using non-linear least squares with the *lmfit* (v0.9.7) package for Python 2.7. When modelling a single class of fitness effects, we estimated *γ_a_* and *p_a_* using Equation 4. To incorporate two discrete classes of advantageous mutational effects, we modified Equation 4, replacing *P_sc,k_* in Equation 4 with

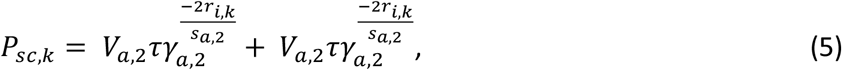

where the subscripts 1 and 2 refer to the two different classes of fitness effects.

To model an exponential distribution of fitness effects, we replaced *P_sc,k_* with

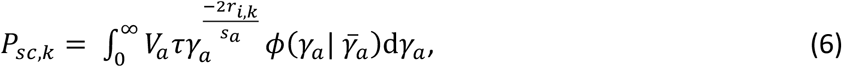

where 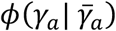 is the probability density function of an exponential distribution with mean 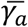. In all cases, we used the average length of protein-coding exons or CNEs, 152.0 and 50.0 respectively, as *τ* when fitting equation 4. We assumed that *N_e_* = 426,200, based on 4*N_e_μ* = 0.0092 and a mutation rate of 5.4 x 10^-9^ (Uchimura et al. 2015).

We used our estimates of positive selection parameters to quantify the relative contributions of positive selection in protein-coding exons versus CNEs to the change in population mean fitness. To obtain confidence intervals around our estimates of the ratio of fitness contributed by positive selection in exons versus CNEs (*ΔW_Exons_/ΔW_CNEs_*), we used a parametric bootstrap approach. For each estimated *γ_a_* and *p_a_* parameter, we sampled random values from a truncated normal distribution with mean equal to the parameter estimate and variance equal to the square of the standard error of the parameter estimate. The truncated normal distribution had a lower bound of 0.0, since values of *γ_a_* and *p_a_* below 0 are biologically impossible. For the calculation of *ΔW_Exons_/ΔW_CNEs_*, we performed 1,000 bootstraps and used them to estimate 95% confidence intervals.

### Data availability

Scripts and code to reproduce all analyses, simulations and figures shown in this study are available at https://github.com/TBooker/MuridRodentProject.

## Author Contributions

TRB, BC and PDK devised the study. TRB, BCJ, RJC analysed the data. TRB wrote the first draft of the manuscript. All authors contributed to the writing and editing of the manuscript.

## Acknowledgements

We would like to extend thanks to Rosina Savisaar for help with identifying exonic splice enhancers and to Peter Ralph for help with PySlim. We would also like to thank Sally Otto, Michael Whitlock, Nathaniel Sharp and Armando Geraldes for helpful discussions, and especially Dan Halligan for help and guidance throughout the course of this and the preceding projects. This project has received funding from the European Research Council under the European Union’s Horizon 2020 research and innovation programme (grant agreement no. 694212).

## Supplementary Material

## Appendix: Analyses assuming LD-based recombination maps

### Generating LD-based recombination rate maps

We phased variant calls using the read-aware methodology incorporated in SHAPEIT2 (Delaneau et al., 2013). For each of the mouse population samples, we carried out the following procedure. First, we created a stringently filtered set of SNPs following Booker et al. (2017), by only including biallelic variant sites that met the following criteria: no overlap with indels, no missing data, QUAL >= 30, genotype quality (GQ) greater than or equal to 15 in all individual genotypes, sequencing depth (DP) greater than or equal to 10 for all individuals, rejected sites with significant deviation from Hardy-Weinberg equilibrium at the level *p* < 0.05). Using the filtered variants, we extracted phase informative reads. We then ran SHAPEIT2 in ‘assemble’ mode to phase our stringently filtered variants. Finally, we converted the output of SHAPEIT2 to FASTA files, which contained two haplotypes per diploid sample using custom Python scripts.

We ran LDhelmet version 1.9 (Chan et al. 2012) on the phased haplotypes, in order to estimate the population-scaled recombination rate, *ρ* = 4*N_e_r*, where *N_e_* is the effective population size and *r* is the rate of crossing over between two sites per generation, for each of the mouse populations. We calculated the ancestral prior probability for each variant site that we passed to LDhelmet using the method developed by Keightley & Jackson (2018) as implemented in the program *est-sfs* v2.01. As input for this program, we generated files including each variant and invariant site that met less stringent filtering criteria than that described above (QUAL > 30, no missing data, no overlap with indels, ExcessHet < 13, no more than two alleles per site), and additionally discarded sites that did not have full outgroup information (alleles from both *Mus famulus* and *Mus pahari* for mouse samples mapped to mm10). For sites that were present in the input to LDhelmet, but not in the input for the Keightley & Jackson (2018) method, because they lacked complete outgroup data, we assigned the ancestral prior following Equation 18 in Keightley & Jackson (2018). We used the resulting information about the ancestral states of SNPs to populate the 4×4 mutation transition matrix used by LDhelmet (Chan et al. 2012). To estimate fine-scale recombination rates in each of our populations, we ran the *find_confs* component of LDhelmet with a window size (-w) of 50 SNPs to generate haplotype configuration files from the phased FASTA files we made in the step above. Subsequently, we ran the *table_gen* and *pade* components of LDhelmet with the default parameters, with the exception of θ [-t] which we set to the point estimate of *π* at 4-fold degenerate synonymous sites specific to each population. To estimate *ρ* we ran the *rjmcmc* component of LDhelmet with a width [-w] of 50 SNPs, a block penalty [-b] of 100, a partition length of 4001 SNPs, an overlap of 200 SNPs, a burn-in period of 100,000 iterations followed by 1,000,000 iterations of the Markov chain.

### Comparison of LD-based recombination rates among taxa

When analysing patterns of genetic diversity under a model of selection at linked sites, the way in which recombination rate estimates were obtained may affect parameter estimates. We analysed the relationship between nucleotide diversity and genetic distance from functional elements in *M. m. castaneus* assuming either a high-resolution recombination map constructed using patterns of linkage disequilibrium (LD) or the pedigree-based map constructed by Cox et al. (2009). These two approaches for generating recombination rate maps have both advantages and disadvantages. By examining patterns of LD, the population-scaled recombination rate (*ρ* = *4N_e_r*), where *r* is the recombination rate, can be inferred from a relatively small sample of unrelated individuals at very fine-scales. However, natural selection can influence LD and may therefore affect such recombination rate estimates (Clark et al. 2010). Alternatively, direct estimates of the recombination rate can be obtained from crossing experiments, but to achieve a high-resolution recombination map, a very large number of individuals need to be genotyped and this has typically precluded the use of whole-genome re-sequencing in some species such as mice, thereby limiting resolution.

We generated recombination rate maps from patterns of LD for each of the mouse taxa, and compared these to the pedigree-based estimates obtained by Cox et al. (2009). It is worth pointing out that the Cox et al. (2009) map is an estimate of the recombination map that was generated using inbred strains of mice of predominantly *M. m. domesticus* origin and there are known differences in total genetic map length and local recombination rate between *M. musculus* sub-species (Dumont & Payseur 2011). For simplicity, we treat the Cox map as a baseline comparison for each of the recombination rate landscapes we inferred.

We calculated Spearman’s correlation between LD-based recombination rate estimates obtained for each mouse taxa and recombination rate estimates from the Cox map in windows from 1Mbp up to 20Mbp. Across all scales tested, the recombination maps for *M. m. castaneus* and *M. m. musculus* from Afghanistan showed the highest level of congruence with the Cox map (Figure A.1). The correlation exhibited by the *M. m. castaneus* was very similar to the correlation previously reported (Booker et al., 2017). For the purposes of calculating genetic distances, we used the LD-based recombination rate estimates for *M. m. castaneus*.

**Figure A.1.**
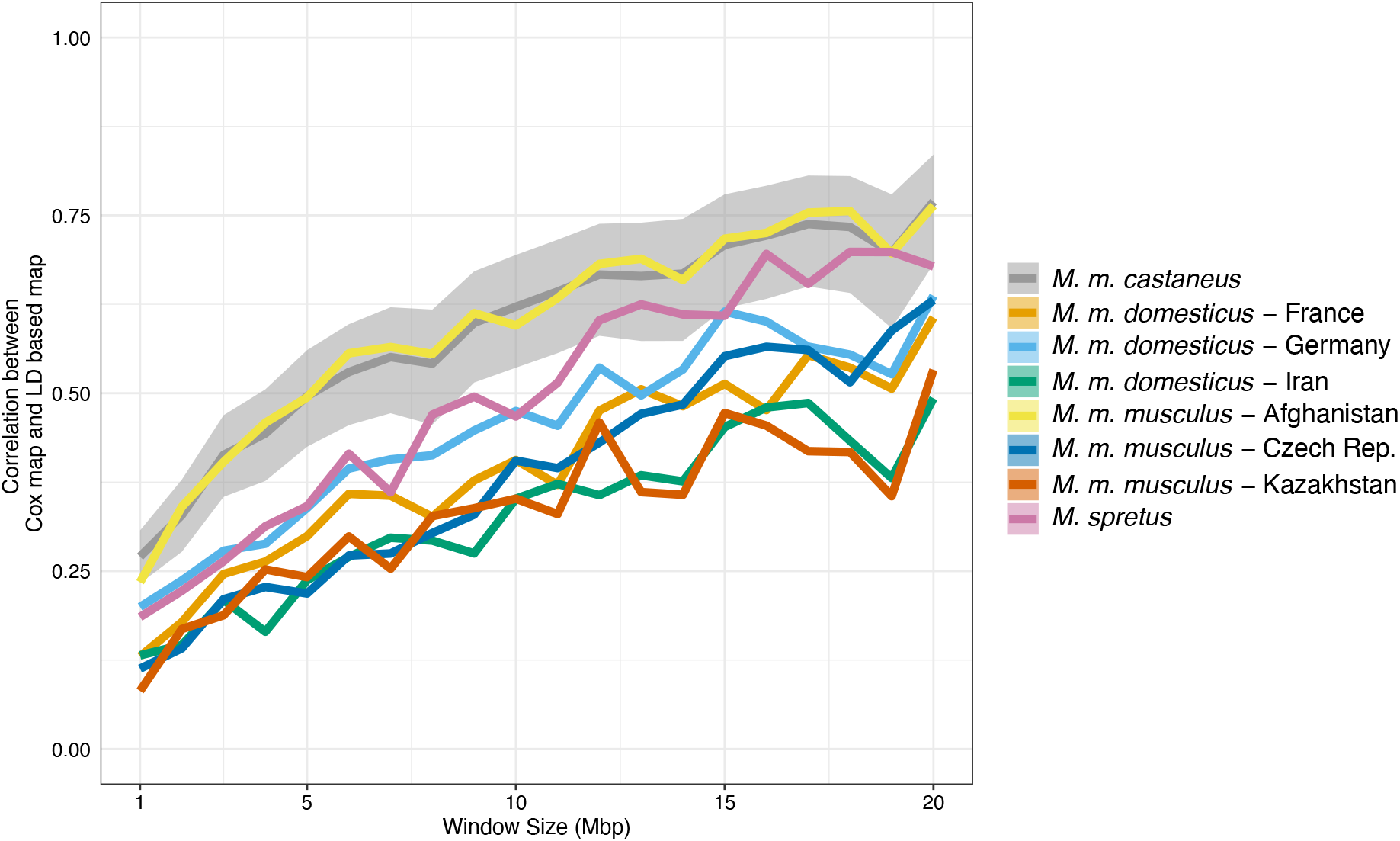
Spearman rank correlation coefficients between recombination maps inferred using LDhelmet for wild mice and the pedigree-based map of Cox et al. (2009). Correlations were calculated in non-overlapping windows of discrete physical size. For the purposes of visualisation, the confidence interval is only shown for the *M. m castaneus* map.

**Figure A.2.**
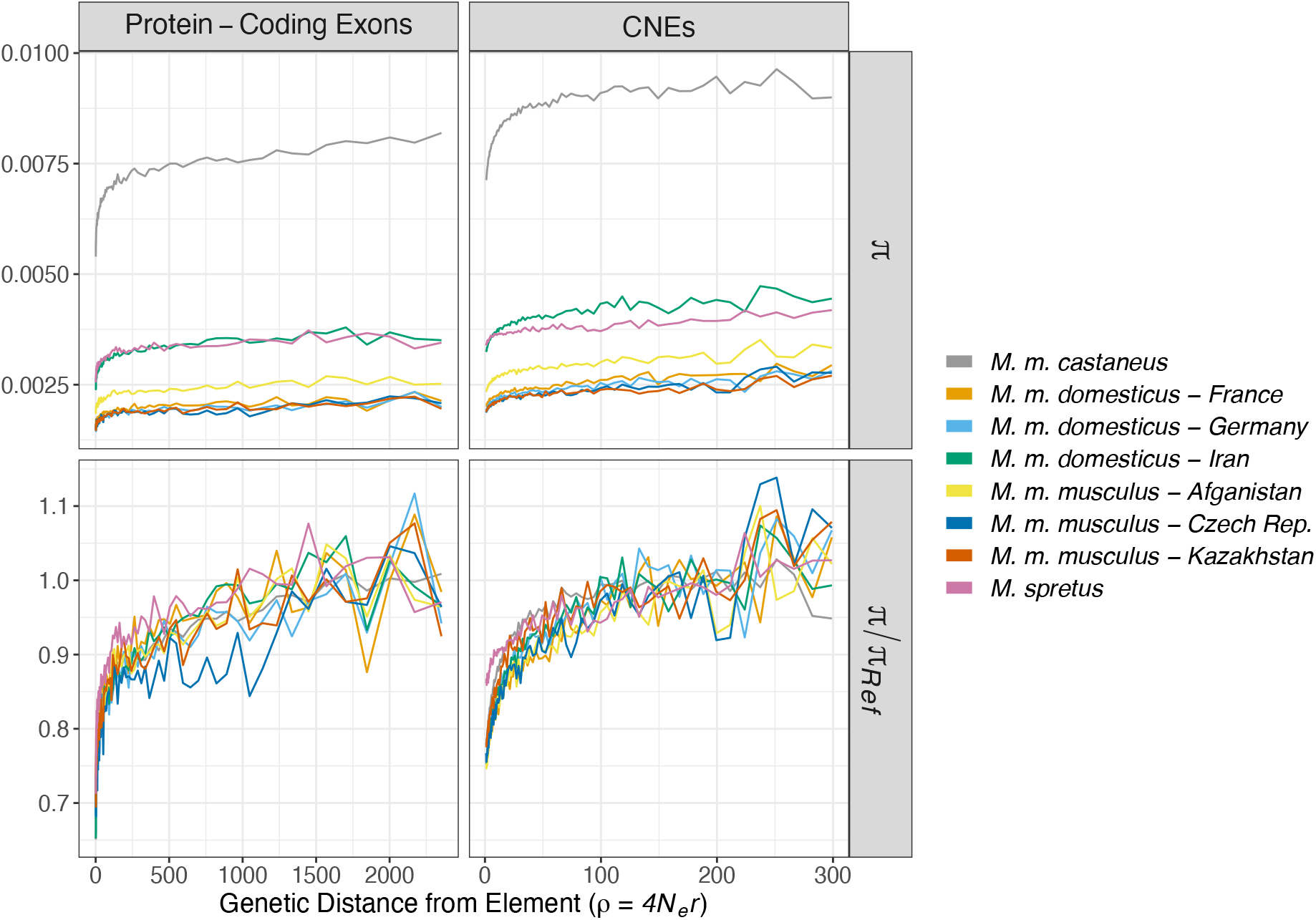
Identical to Figure 2 in the main text except that genetic distances were calculated assuming the LD-based recombination map constructed for *M. m. castaneus*.

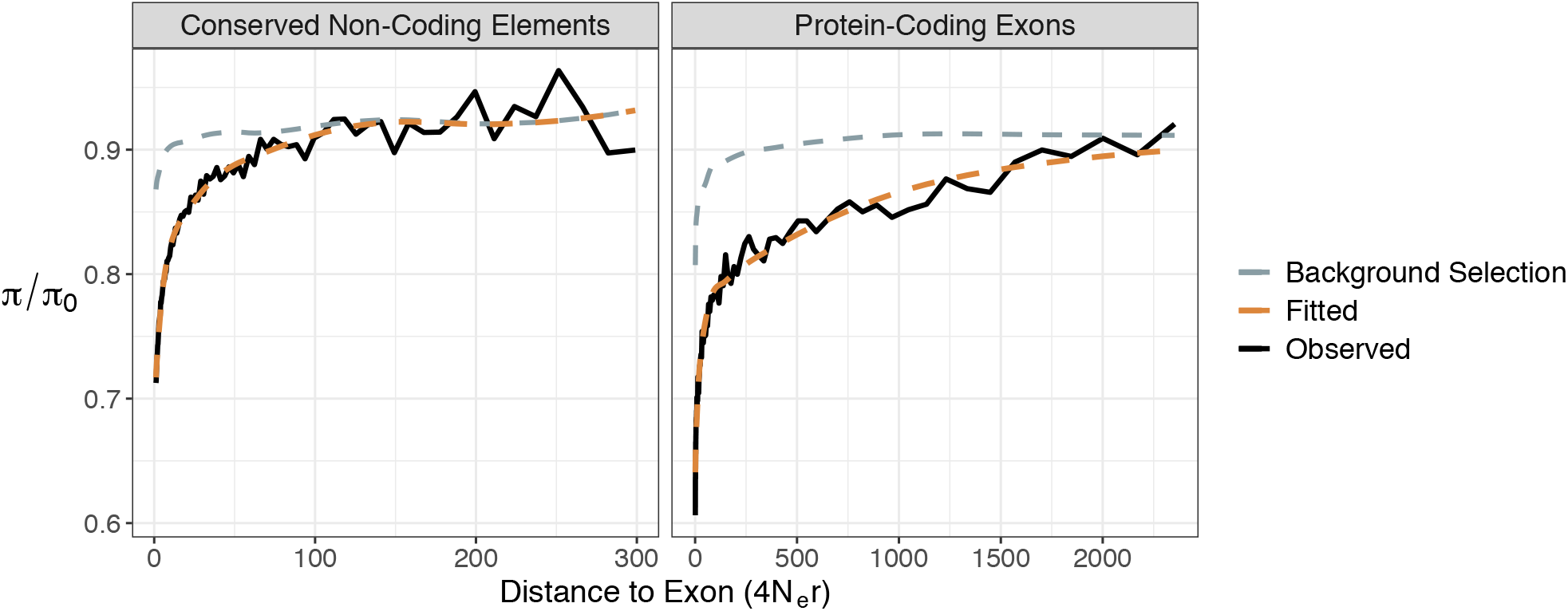

**Supplementary Table S1** Comparison of uSFS model fits for each taxa and class of sites considered. The maximum likelihood estimate of model parameters are shown along with the estimated uSFS. A parameter key is given as a second sheet in the spreadsheet.

**Supplementary Table S2** Parameters of the distribution of fitness effects for deleterious mutations as well as the positive selection parameters estimated for each population using *polyDFE*. Point estimates are provided as well as 95% bootstrap confidence intervals. A parameter key is given as an additional sheet in the spreadsheet.

**Supplementary Table S3** Estimates of positive selection parameters obtained by fitting a models of selective sweeps and background selection to troughs in nucleotide diversity. Parameters are given for models assuming a one or two discrete classes of advantageous mutations as well as an exponential distribution of fitness effects. Estimates of the fitness change brought about by positive selection in protein-coding exons and CNEs are also given in the table. A parameter key is given as an additional sheet in the spreadsheet.

**Figure S1.**
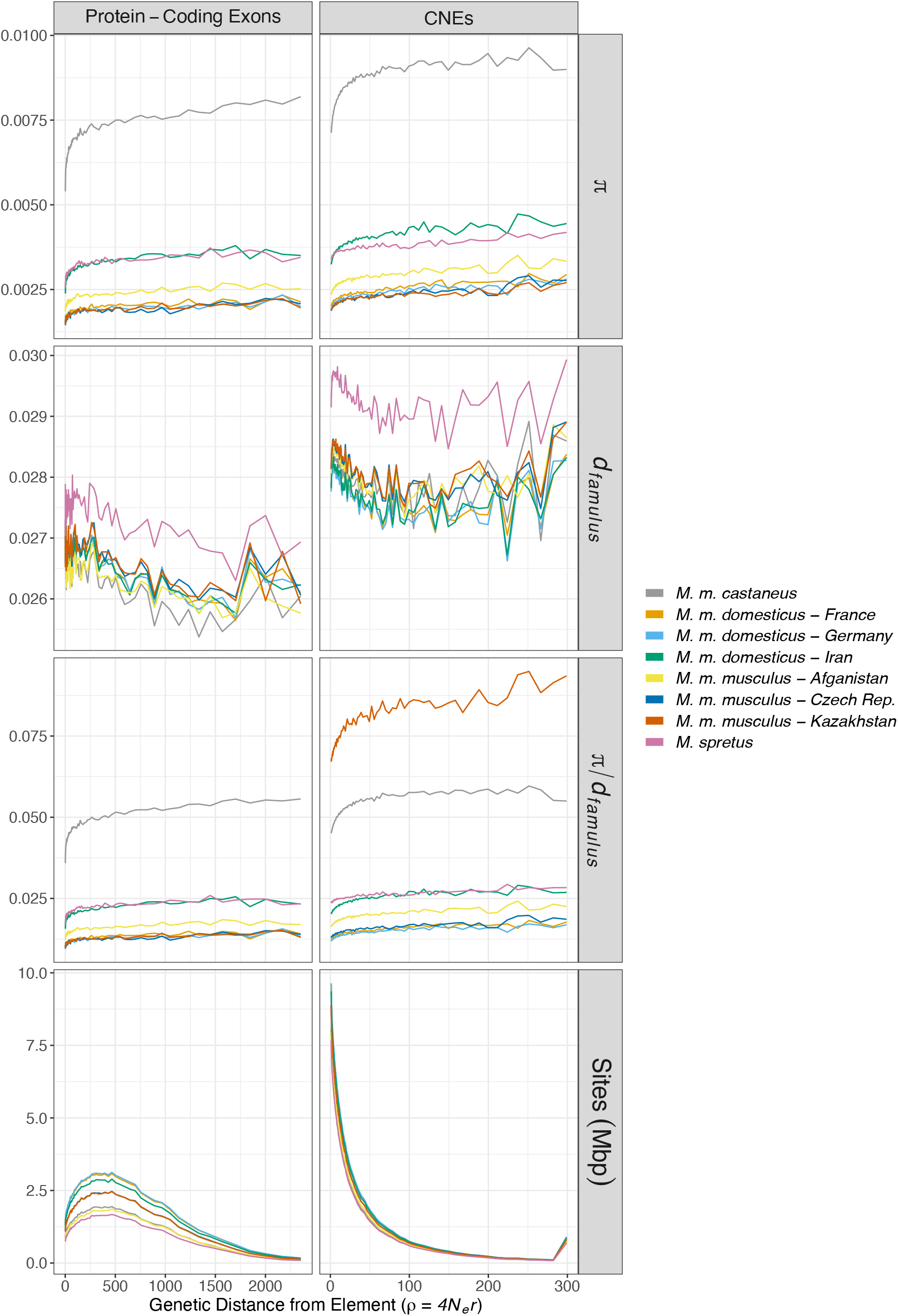
Additional summary statistics in the regions surrounding functional elements assuming the LD-based map of recombination rate variation we inferred for *M. m. castaneus*.

**Figure S2.**
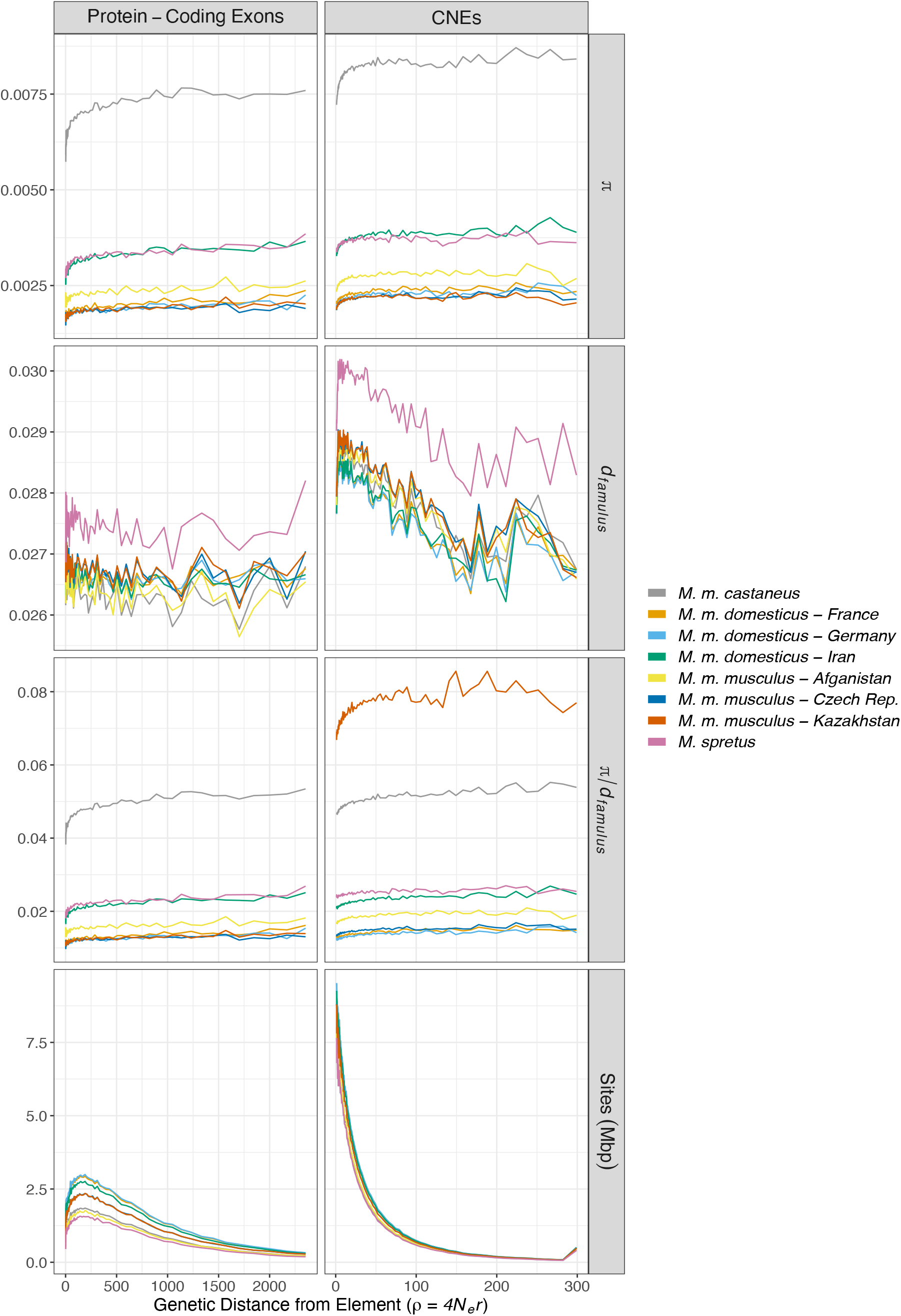
Additional summary statistics in the regions flanking functional elements assuming the Cox map.

**Figure S3.**
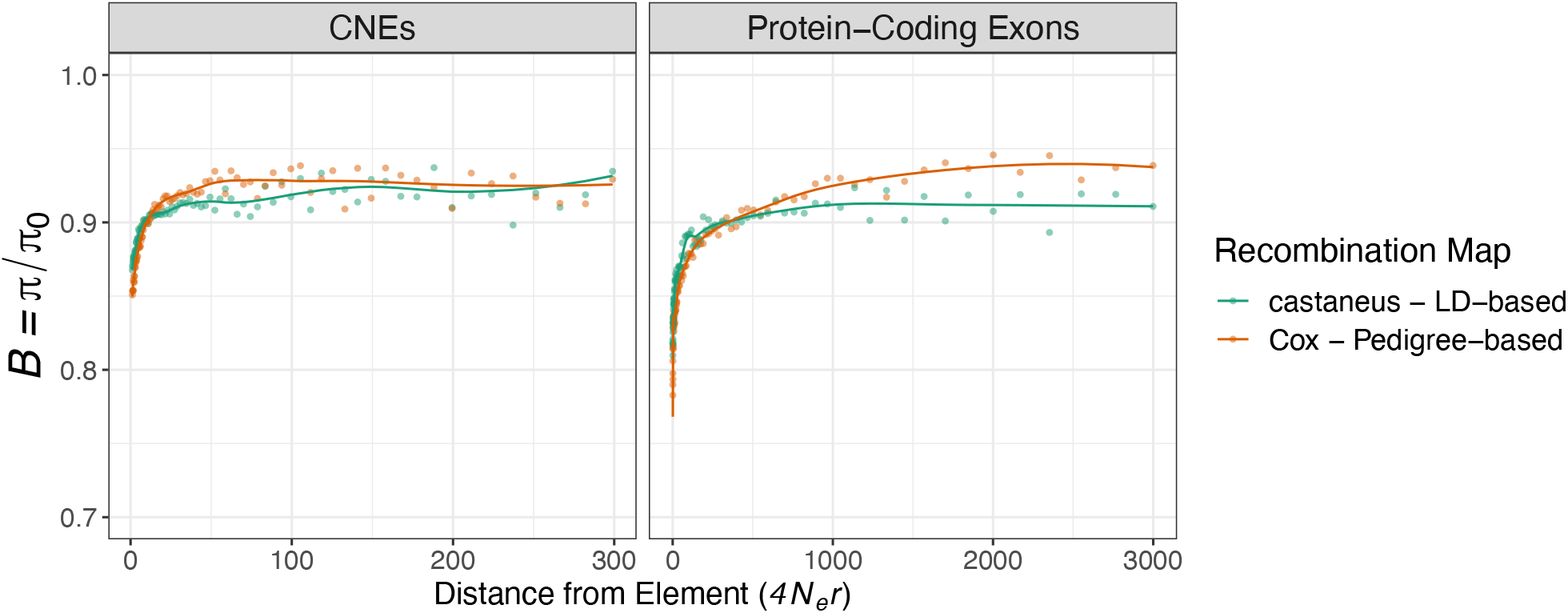
The reduction in neutral genetic diversity relative to neutral expectation caused by background selection (*B*) observed in simulated datasets. Simulations assumed either the LD-based recombination map or the pedigree-based map of Cox et al. (2009). Lines indicate the fit of a Loess regression fitted to the data with a span of 0.3 and the number of sites in each bin used as weights.

**Figure S4.**
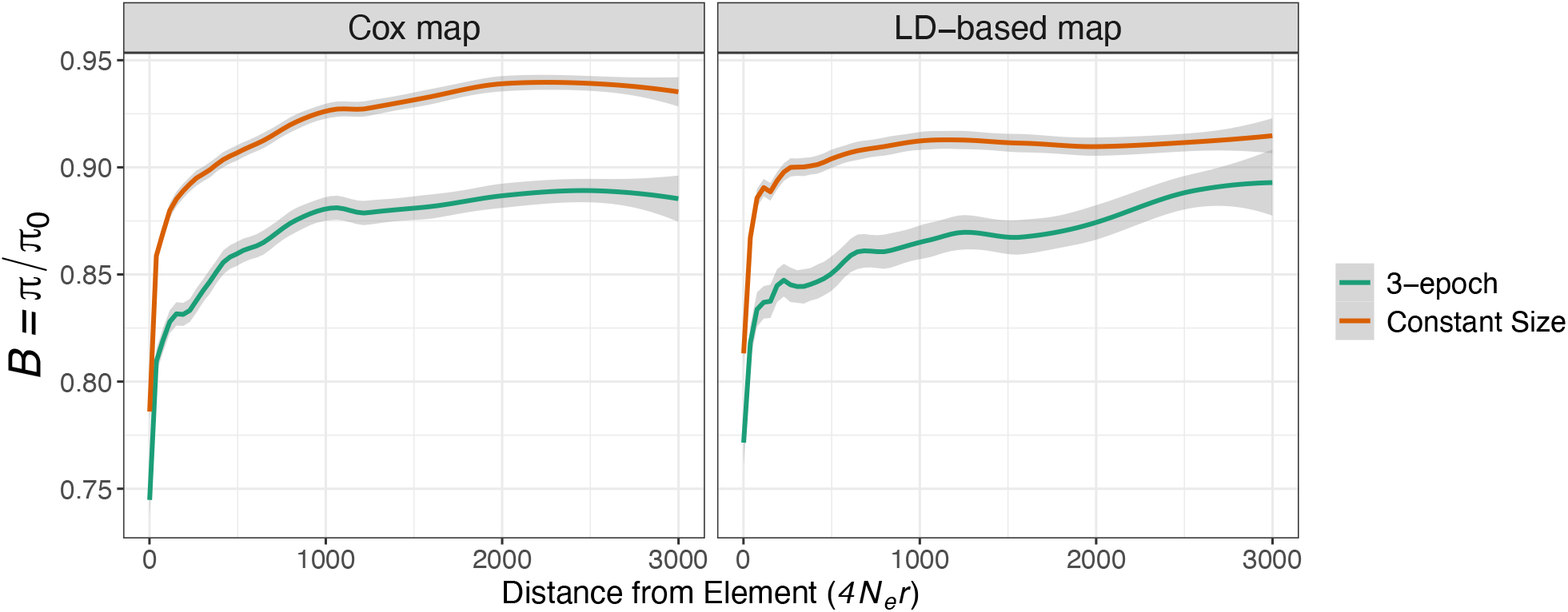
The reductions in neutral genetic diversity relative to neutral expectation caused by background selection (*B*) observed in simulated datasets when modelling a population with constant size, or the three-epoch demographic model estimated by Booker and Keightley (2018). π_0_ in the constant size simulations was 0.01. π_0_ was 0.0042 in the 3-epoch simulations, which was calculated from the harmonic mean of population sizes. Lines indicate the fit of a Loess regression fitted to the data.

